# Identification of Antibiofilm Agents against *Salmonella enterica* from the Pathogen Box Compound Library

**DOI:** 10.64898/2026.02.26.707623

**Authors:** Adeshola A. Fagbemi, Chinedum P. Babalola, David A. Kwasi, Olabisi C Akinlabi, Olayinka Kotila, Iruka N. Okeke

## Abstract

**Background:** Biofilms are central to *Salmonella* pathogenesis, and targeting their formation is believed to produce less evolutionary pressure of growth inhibition than traditional antibacterials. In this study, we screened the Medicines for Malaria Venture (MMV) Pathogen Box library to identify anti-biofilm agents against *S*. *enterica* that possess drug-like properties.

**Methodology/ Principal Findings:** A crystal-violet-based medium-throughput antibiofilm screen of *Salmonella enterica* serovar Typhimurium ATCC 14028 and a clinical *Salmonella enterica* serovar Elisabethville isolate was performed on polystyrene surfaces using the 400-compound Pathogen Box library. Compounds that inhibited biofilm formation by >30% and growth by <10% were identified as hits. *Salmonella* red-dry-rough and motility phenotypes were explored in mechanism of action studies on one hit compound. The *Salmonella* antibiofilm hit rate was 0.75% for this library. MMV688371 (benzamide) inhibited biofilm formation of *S*. Typhimurium ATCC 14028 by 33% without inhibiting growth. An ethambutol analogue (MMV687273) and auranofin (MMV688978) met the hit criteria against *S*. Elisabethville LLD035X. Auranofin showed concentration-dependent, growth-inhibition-independent antibiofilm activity against typhoidal and non-typhoidal *Salmonella* from Nigeria, and inhibited the motility of *S*. Elisabethville LLD035X at 5 µM. At 5 µM aurothioglucose, an auranofin gold (I) analogue, and non-gold analogue 1-Thio-beta-D-glucose tetraacetate, inhibited biofilm formation by 61.30% and 11.39%, respectively, pointing to essentiality of the gold (I) moiety for activity.

**Conclusions/ Significance:** Structurally diverse small molecules can inhibit biofilm formation by *Salmonella,* and motility inhibition is an important mechanism for this activity. Auranofin inhibits typhoidal and non-typhoidal *Salmonella* biofilm formation, with its gold content being required for these activities.

## Background

*Salmonella enterica* serovars are widely disseminated and high-burden foodborne and waterborne pathogens that can survive in abiotic and biotic environments (Yada, 2023; Mkangara, 2023). Biofilms, which are cellular and polymer matrix formations that serve as an enclosure and reduce or eliminate the effect of external stressors on these organisms, contribute to *Salmonella* pathogenesis and survival (Aleksandrowicz et al., 2023) as well as to antimicrobial resistance and treatment failure (Moshiri et al., 2018). Biofilm structural components include live cells, extracellular DNA, glycolipids, glycoproteins, and peptidoglycan-linked polysaccharides. Surface attachment, auto-association, and matrix encapsulation processes that are necessary for biofilm formation support the recurrence of infection caused by *Salmonella enterica* by prolonging microbes’ sustainability and release, aiding survival under host and environmental constraints, and increasing the probability to colonize a new host (Koopman et al., 2015; Xiao et al., 2017).

*Salmonella enterica* subspecies enterica consists of serovarieties that are typhoidal (Typhi and Paratyphi) and non-typhoidal. Typhoidal serovars cause typhoid or paratyphoid fever and are human-specialized pathogens. Non-typhoidal *Salmonellae* are zoonotic and mainly causative agents of gastroenteritis, although bacteriemia and invasive infections are occasionally attributed to them, and a few lineages, notably *Salmonella enterica* serovar Typhimurium ST313 and *Salmonella enterica* serovar Enteritidis ST11, commonly cause invasive disease. Invasive non-typhoidal *Salmonella* disease annual incidence in sub-Saharan Africa exceeds 100 per 100,000 individuals, in endemic settings (114 per 100,000 person-years of observation (95% CI 91–140) in Nigeria) with 16% (46 of 280) of *S*. Typhi isolates tested showing ciprofloxacin non-susceptibility (Marks et al. 2024). Biofilm formation plays a crucial role in the persistence, dissemination, development of resistance, and survival of *Salmonella*, the discovery of compounds capable of inhibiting biofilm development represents a critical and promising avenue for research and therapeutic intervention.

Small molecules have been shown to inhibit biofilm formation of a wide range of organisms, many acting on surface factors or their regulation (Kalia et al., 2023). Pathogen Box, compiled and distributed by Medicine for Malaria Venture (MMV) is a curated chemical library containing 400 structurally diverse but druggable chemical compounds reported to exhibit biological activity and to be less cytotoxic to mammalian cell lines. We hypothesised that this box contains antibiofilm scaffolds active against *Salmonella enterica* and screened them in medium throughput library.

## Methods

### Bacterial Strains and Growth Conditions

This study used *Salmonella* Typhimurium strain ATCC 14028 and clinical isolates (Table 1) obtained from salmonellosis surveillance in Nigeria. Bacterial stocks were preserved in Luria-Bertani (LB) broth: glycerol (1:1) at −80°C. For experimentation, bacteria were routinely cultured on Tryptic Soy Agar or in Luria-Bertani broth with shaking at 150 rpm, at 37°C.

**Table 1:**
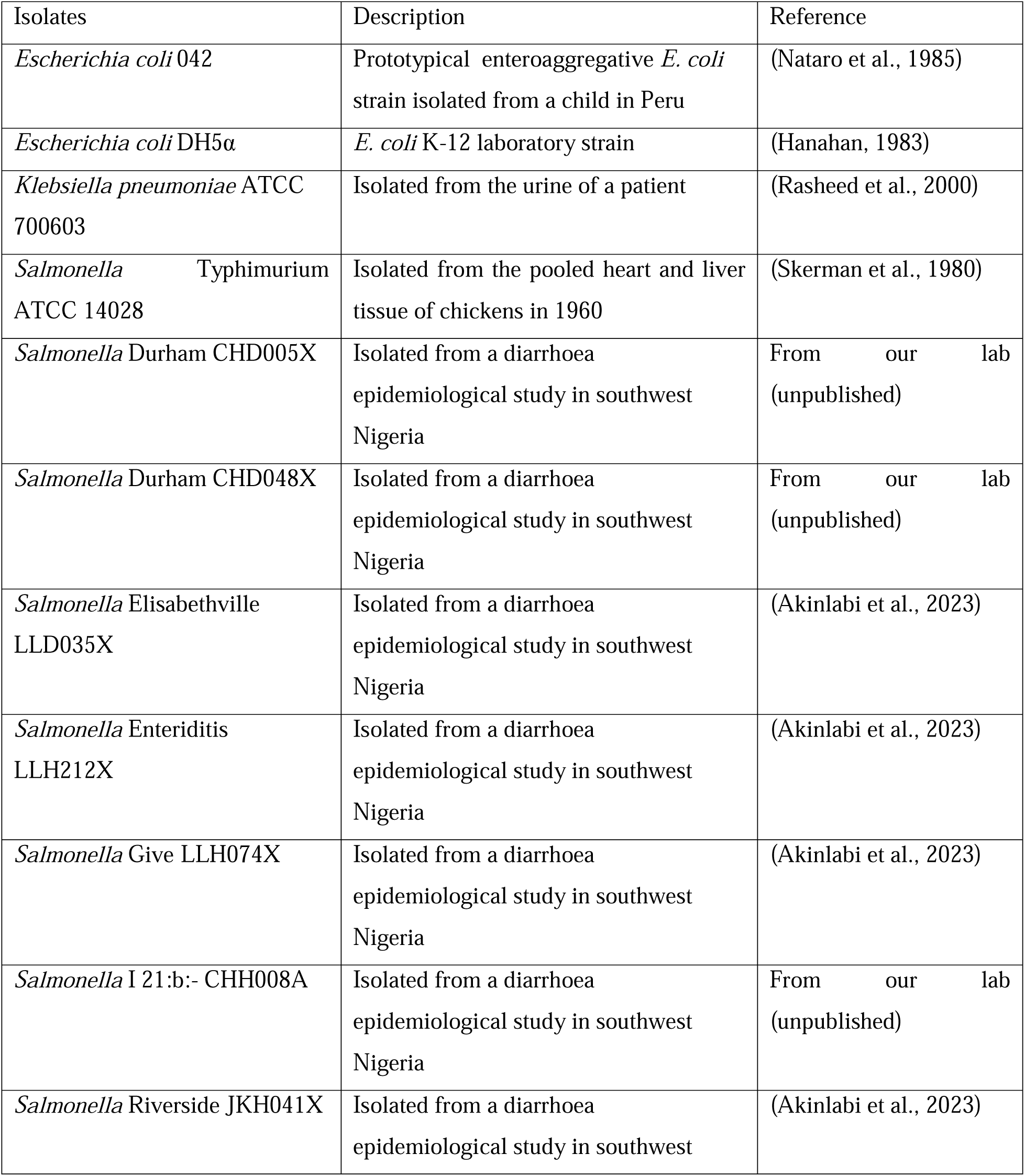
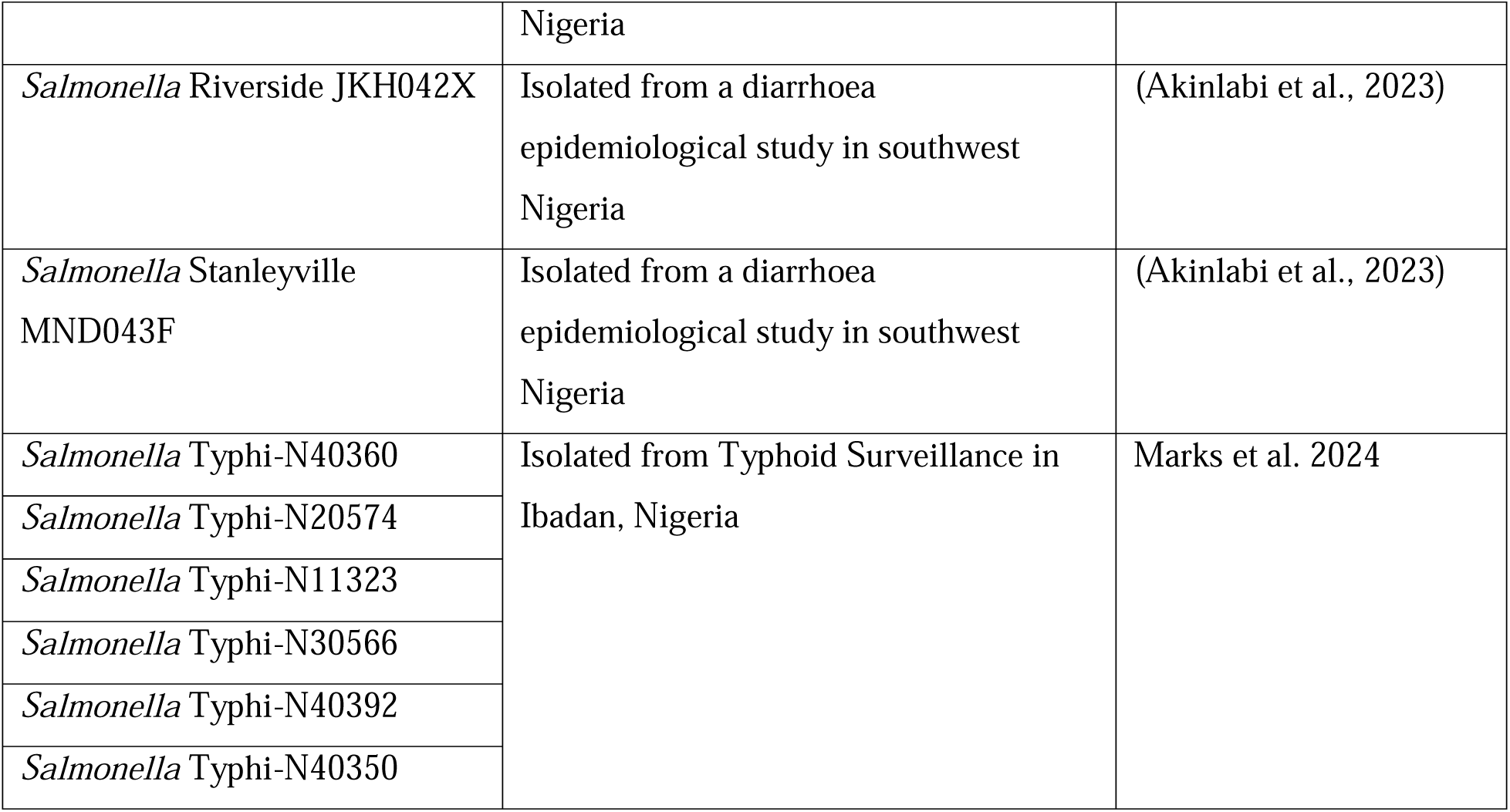
Bacterial strains used for this study.

**Table 2:**
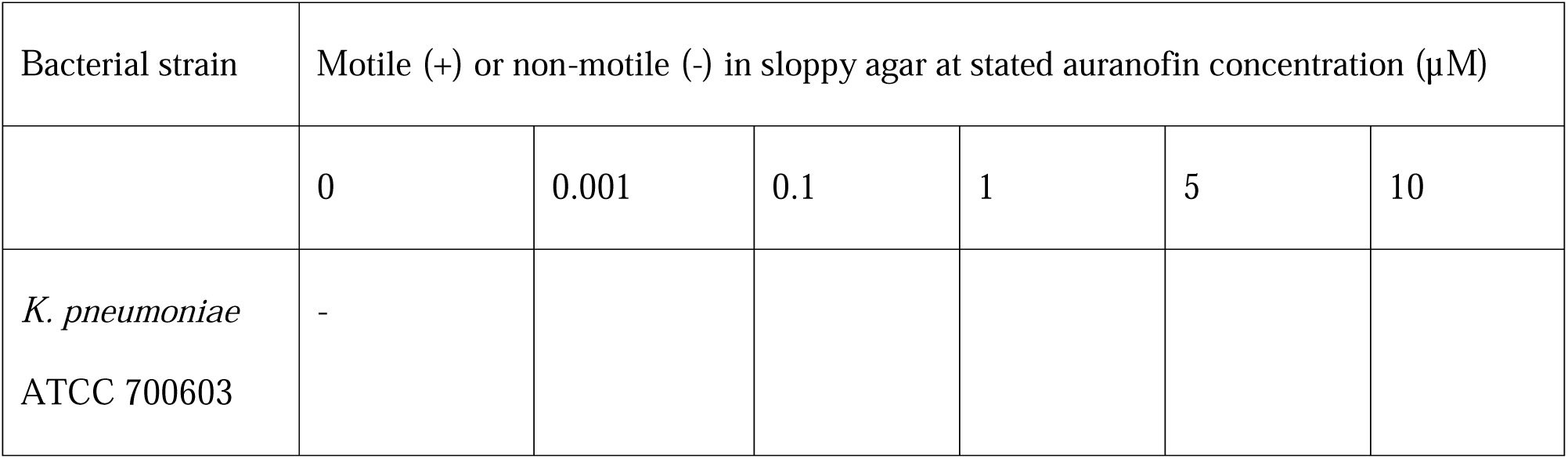
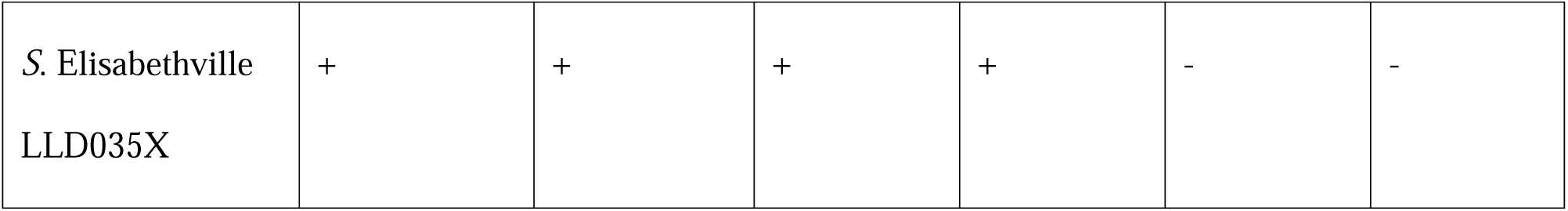
MMV688978 inhibition of *Salmonella* swimming motility on soft agar medium.

### Test compounds

The Pathogen Box library was gratefully received from Medicine for Malaria Venture (MMV). The library held 400 predominantly structurally distinct molecules, with a limited number of related structures also present. Compounds returned as active from the initial antibiofilm screening were requested from the Medicine for Malaria Venture for further investigation and MMV675968, MMV688179 and MMV687747 were supplied. Auranofin (Sigma-Aldrich), aurothioglucose, and 1-thio-beta-D-glucose tetraacetate were purchased from MedChemExpress. Compounds were dissolved in DMSO and tested in DMSO: sterile distilled water at a 1:2 ratio.

### Evaluation of strain biofilm formation in different media

Biofilm formation by *Salmonella* Durham CHD005X, *Salmonella* Durham CHD048X, *Salmonella* I 21:b:-CHH008A, *Salmonella* Elisabethville LLD035X, *Salmonella* Riverside JKH041X, *Salmonella* Riverside JKH042X, *Salmonella* Give LLH074X, *Salmonella* Give LLH072X, *Salmonella* Enteritidis LLH212X, *Salmonella* Typhimurium ATCC 14028, *Salmonella* Stanleyville MND043F was tested in Dulbecco’s Modified Eagle Medium (DMEM), Luria-Bertani broth, Tryptic Soy Broth, Tryptic Soy Broth+ 2.5% glucose, 0.05 × Tryptic Soy Broth at 37□ at static condition and 28□ at 50 rpm. Enterobacterales of known biofilm forming capacity such as enteroaggregative *Escherichia coli* 042 (Sheikh et al., 2001), and *Klebsiella pneumoniae* ATCC 700603 (Shadkam et al., 2021) and *Escherichia coli* DH5α (K-12 laboratory strain and non-biofilm former) were used as controls.

### Antibiofilm screen

The medium-throughput screen of the library against *S*. Typhimurium ATCC 14028 and *S*. Elisabethville LLD035X was performed as described by Kwasi et al (2022), with slight modifications. The screen used sterile polystyrene flat-bottom plates (96-well). Each well of these plates was set up with 5 μL of the stock compound solution, bacteria (grown in broth and diluted 1:10 in a 1:20 dilution of Tryptic Soy Broth, or TSB), and additional medium (1:20 TSB) to reach a final volume of 200 μL, resulting in a chemical concentration of 5 μM of the compound. For *S*. Typhi and *Klebsiella pneumoniae* ATCC 700603, TSB was used in place of 1:20 TSB. Each plate had a control vehicle of 5 μL of DMSO: H_2_O (1:2) made up to 200 μL of medium. The assay plates were prepared in triplicate and incubated at 28°C with shaking at 50 rpm for 24 hours. Afterward, planktonic cell growth was measured by optical density at 595 nm using a Multi-skan microplate reader. To quantify biofilm formation, the non-adherent cells were carefully aspirated, plates were subjected to three washes with phosphate-buffered saline using an ELISA microplate washer, and then dried. Biofilms were fixed by 75% adding ethanol and standing for 10 minutes, after which the plates were dried again. 200 μL of 0.5% crystal violet was added to each well for 5 minutes without agitation. The unattached dye was discarded, and plates were thoroughly washed, and dried. The crystal violet stain in the biofilm was eluted by adding 200 μL of ethanol at 95% strength to each well, and 150 μL of the resulting solution was introduced to a new corresponding well in a fresh plate for quantification. The biofilm formed was measured by the optical density of this solution at 570 nm using a Multi-skan microplate reader. Hits were selected based on antibiofilm activity greater than 30%, compared to wells without added compound, and growth inhibition of less than 10%.

To validate hits from the antibiofilm screen, biofilm formation over a concentration gradient of 0–10 μM and varying time intervals, 6–48 h was tested, using the afore-described setup. The antibiofilm spectra of hit compounds were determined by testing a range of *Salmonella* and other species (Table 1) against hit compounds.

### Phylogenetic mapping

Reads were aligned to a reference genome (SRR35837553), and SNPs called out from the multi-fasta alignment generated using snp-sites (https://github.com/satta/snp_sites). A SNPs phylogenetic tree was then constructed with IQ-TREE www.iqtree.org (Minh et al., 2020) using GTR+G model and visualized with Microreact https://microreact.org.

### Bacterial motility test in sloppy agar

To evaluate the impact of MMV688978 on *Salmonella* motility, swimming motility assay on soft agar medium was deployed. A colony of the tested strains were stabbed into 0.3% semisolid agar medium in a 24 well plate. Following the application of gradient concentrations of the MMV688978 (from 0 to 10 μM), the plates were incubated. Motility was quantified by measuring the diameter of motility from the point of inoculation after 18 h incubation at 37 °C.

### Bacterial curli morphotype

Bacterial strains were streaked on Congo red plates (1% casamino acids, 0.1% yeast extract, 20 µg/ml Congo red and 10 µg/ml Coomassie brilliant blue G) and curli expression was monitored for 48 h at 28°C.

### Statistical analysis

Data were statistically analysed using R Studio Software. Data are presented as inhibition, percentage inhibition, and means ± standard error of the mean of replicate readings or median as appropriate, the concentration dependent assay were analysed by one-way ANOVA followed by Dunnett’s multiple comparisons using R Studio Software. P values of <0.05 was considered significant.

## Results

### *Salmonella enterica* form robust and reproducible biofilms in tryptic soy broth (TSB)

The control strains and local strains used in this study are listed in Table 1. To identify *Salmonella enterica* strains capable of forming biofilms, and the optimal conditions for this process, a preliminary study was conducted with a control strain known to form biofilms (*E. coli* 042) and a strain incapable of robust biofilm formation (*E. coli* DH5α). The strains tested included *S*. Typhimurium ATCC 14028, as well as clinical Salmonella isolates. Based on reports from existing literature (Paytubi et al., 2017; Römling et al., 2000; Zogaj et al., 2001), four different media (Dulbecco’s Modified Eagle Medium (DMEM), 1X Tryptic Soy Broth, 0.05XTryptic Soy Broth (1:20 TSB), and Tryptic Soy Broth + 2.5% glucose) and two incubation temperatures (28°C and 37°C) were evaluated to determine the optimal biofilm formation conditions for the *Salmonella*. Control strain enteroaggregative *E*. *coli* 042, formed its strongest biofilms in DMEM at 37°C, as previously reported by Sheikh et al (2001), and laboratory *E. coli s*train DH5α did not form significant biofilms under any of the test conditions used. TSB media supported biofilm formation in most of the *Salmonella* isolates, 1:20 TSB at 28°C was employed for the preliminary antibiofilm screening, specifically, because it represents the environmental stress/starvation state where ex-vivo biofilm-specific genes are most active (Castelijn et al., 2012). The wild-type strain *S*. Typhimurium ATCC 14028 and a local strain, *S*. Elisabethville LLD035X, isolated from a study of infants with diarrhoea (Akinlabi et al, 2023), and were selected for the initial biofilm inhibition screening.

### Pathogen Box contains compounds that inhibit formation of *S*. Elisabethville LLD035X and *S*. Typhimurium ATCC biofilms

Four hundred compounds from the Pathogen Box were screened at 5 μM to determine whether they could prevent or slow biofilm formation by *S*. Elisabethville LLD035X and *S.* Typhimurium ATCC 14028 1:20 TSB at 28°C. The outcome of the screen, performed in triplicate, is depicted in Figure 2(a - c), The majority of the compounds in Pathogen Box did not inhibit the growth or biofilm formation by either strain, and a total of 202 and 321 compounds, against *S.* Typhimurium ATCC 14028 and *S.* Elisabethville LLD035X, respectively (including all three hit compounds), actually promoted bacterial growth, that is recorded ‘growth inhibition’ of <0%. Levofloxacin (MMV687798), amikacin (MMV687796), rifampicin (MMV688775), doxycycline (MMV000011) are antibacterial compounds in Pathogen Box with known Gram-negative inhibitory activity: they exhibited percentage biofilm inhibition and percentage growth inhibition of 77.22% and 50.95%, 80.61% and 30.53%, 70.15% and 42.29%, and 46.92% and 36.70%; respectively against *S.* Elisabethville LLD035X (Figure 2 b and c). For *S.* Typhimurium ATCC 14028, their percentage biofilm inhibition and percentage growth inhibition are levofloxacin (82.50% and 27.44%), amikacin (11.21% and 21.45%), rifampicin (79.09% and 69.34%), and doxycycline (68.05% and 43.58%). At a concentration of 5 μM, MMV687273 and MMV688978 met hit criteria by inhibiting *S*. Elisabethville LLD035X biofilm formation relative to the control (DMSO solvent) by 41.18% (p= 0.0039) and 62.88% (p = 0.0003), respectively. Bacterial growth was not affected by these compounds at 5 μM, with MMV687273 showing an inhibition of −6.13% (p= 0.1398) and MMV688978 showing an inhibition of −14.35% (p= 0.0037), as illustrated in Figure 2 and Figure 3 (Negative inhibitory values indicate that the compound supported growth). Compounds MMV687273 and MMV688978 did not meet hit criteria against inhibited S. Typhimurium ATCC 14028 because they only inhibited biofilm formation by 27.49% (p = 0.006) and 60.60% (p < 0.0001), respectively, but MMV688978 inhibited growth by 17.80% (p = 0.036). Similarly, at 5 μM, MMV688371 inhibited *S*. Typhimurium ATCC 14028 biofilm formation and growth by 33.36% (p = 0.0150) and −17.61% (p = 0.0032), respectively. MMV688371 inhibited biofilm formation and growth of *S*. Elisabethville LLD035X by 8.90% (p = 0.35) and −19.00% (p = 0.11), respectively. Our preliminary medium-throughput screens identified some other potential candidates, including MMV675968 (indicated for cryptosporidiosis) as a hit against *S*. Elisabethville LLD035X, as well as MMV688179 (a kinetoplastid agent) and MMV687747 (an antitubercular agent) against *S*. Typhimurium ATCC 14028. However, on subsequent assay to validate the hits via concentration-dependent assays, they did not demonstrate antibiofilm activity that met the hit criteria. Consequently, these compounds were excluded from the list of hits.

**Figure 1.**
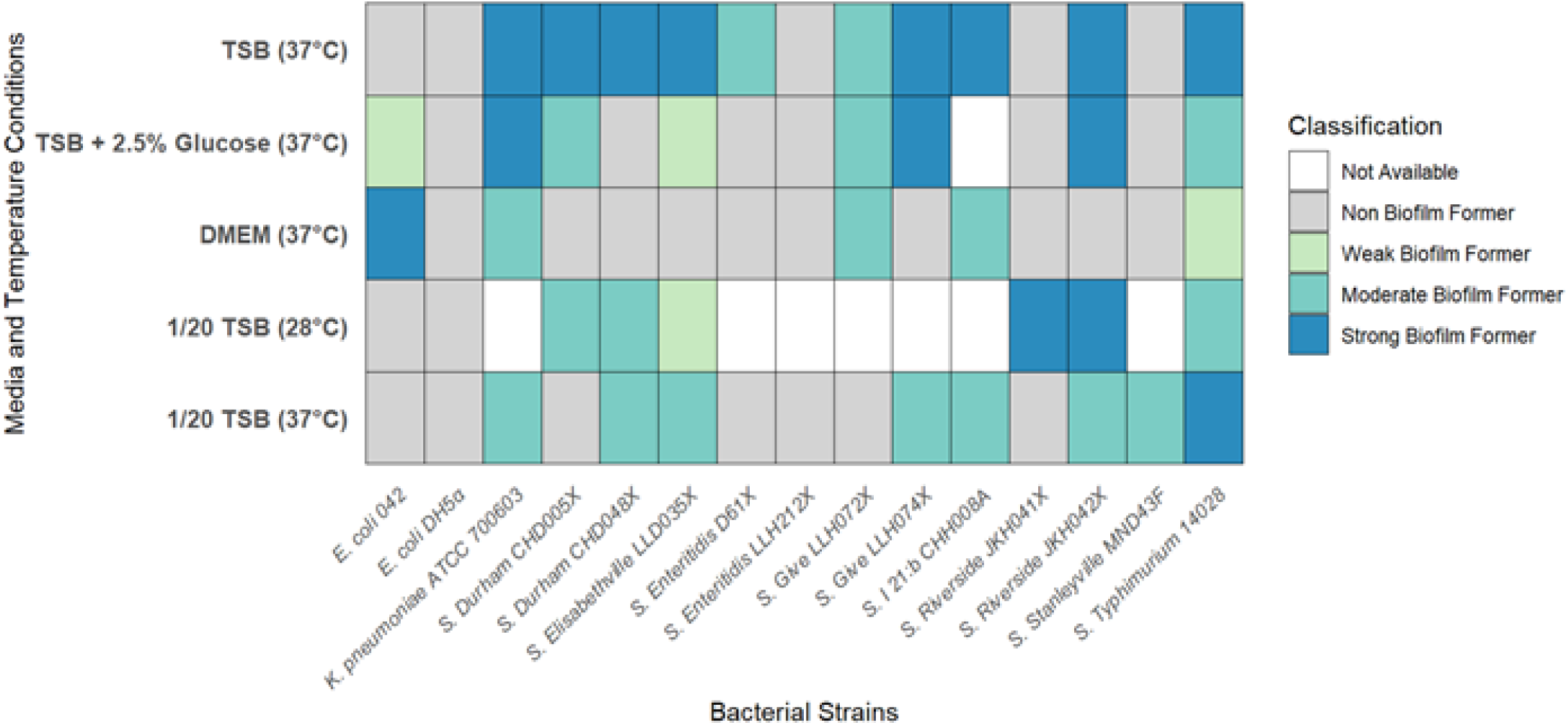
Preliminary biofilm assays conducted to identify biofilm-producing strains and the culture media and temperature conditions that promote biofilm formation. Biofilm formation was determined at 570 nm. Bacteria are classified into four distinct groups based on their ability to form biofilm. Strong Biofilm Former are represented as dark blue, Moderate Biofilm Former are coloured teal, Weak Biofilm Former are shaded light green and Non Biofilm Former are represented as grey. Samples coloured white are missing data i.e., not available. Classification is based on comparing the Optical Density of the sample to an Optical Density of the negative control (*E. coli* DH5α).

**Figure 2.**
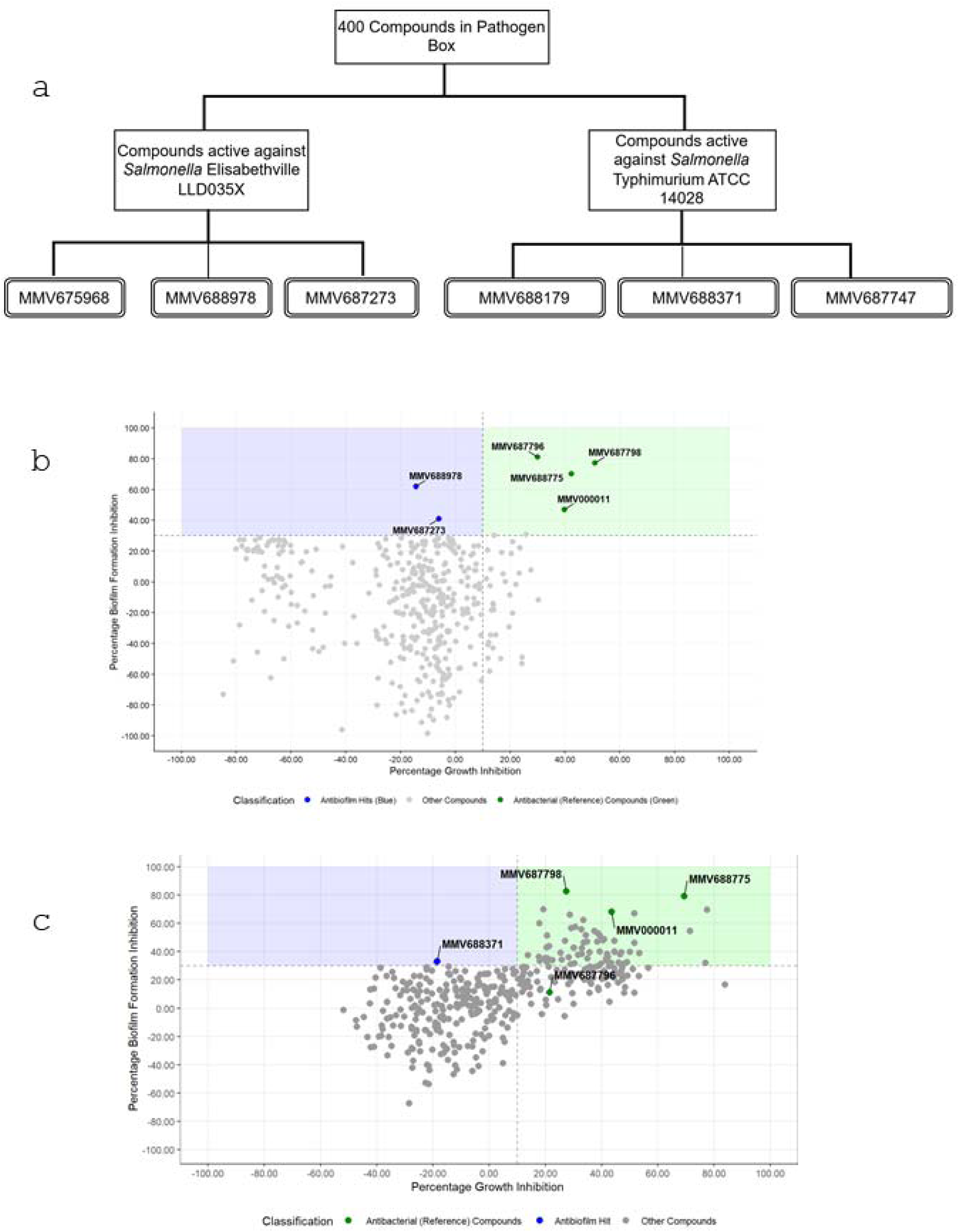
Medium-throughput screen of the Pathogen Box library against *S*. Elisabethville LLD035X and *S*. Typhimurium ATCC 14028 biofilm formation (a-c). (a) Overview of the hit cascade used to prioritize compounds, resulting in two biofilm displaying none or minimal growth-inhibitory hits against *S*. Elisabethville LLD035X and one hit against *S*. Typhimurium ATCC 14028. MMV675968, MMV688179 and MMV687747 were initially picked as hits but were removed later because they could not reproduce antibiofilm activity. MMV687273 and MMV688371 were not available for further investigation. (b) & (c) Preliminary screening of 400 Pathogen Box compounds showing percentage biofilm inhibition versus percentage growth inhibition at 5 μM for *S*. Elisabethville LLD035X (b) and *S*. Typhimurium ATCC 14028 (c). The hit compounds are represented as blue and fall within the hit zone shaded as light blue. Compounds in Pathogen Box with known antibacterial effect against Gram-negative bacteria are represented in emerald green while the other non-hit compounds are shown in slate grey. Compounds above the horizontal line exhibit >30% biofilm inhibition, while those left of the vertical line show <10% growth inhibition. Two hits for *S*. Elisabethville and one hit for *S*. Typhimurium were identified.

**Figure 3.**
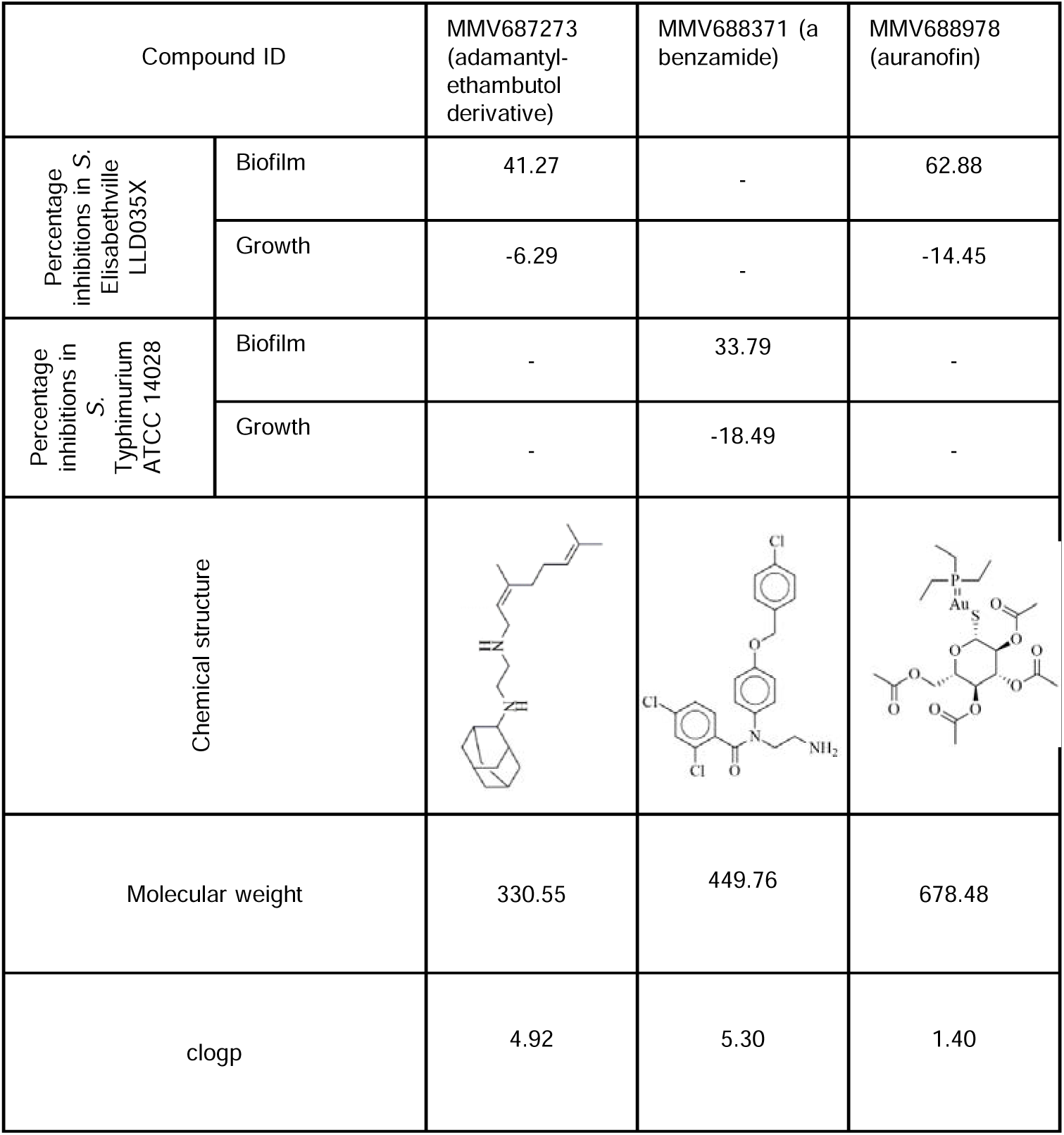
Chemical structures and key molecular descriptors of hit compounds active against *S*. Elisabethville LLD035X and *S*. Typhimurium ATCC 14028, with data provided by MMV.

Thus, three compounds met the hit selection criteria of ≥30% inhibition of biofilm formation and ≤10% inhibition of growth, MMV687273 and MMV688978 were identified as hits against *S*. Elisabethville LLD035X (hit rate = 0.50%) while MMV688371 was the active compounds for *S*. Typhimurium ATCC 14028 (hit rate = 0.25%). No compound met hit criteria against both strains. As depicted in Figure 3, in addition to not having overlapping spectra, the compounds were structurally unrelated. The hits MMV687273 and MMV688371 were not available commercially. We therefore focused on validating and characterizing MMV688978, which is known as auranofin.

### Auranofin (MMV688978) demonstrates dose-dependent inhibition of S. Elisabethville LLD035X biofilm formation and no significant effect on growth

Following established protocols, strains were incubated for biofilm formation for up to 24 hours in the presence of MMV688978 at concentrations ranging from 0.00 µM to 15.00 µM. Biofilm formation was inhibited in a dose-dependent manner, optimum inhibition of 72.08% (P < 0.001) was seen at 1.25 µM and 15.00 µM (Figure 4(a)). Before crystal violet quantitation of the biofilms, plates were read at OD_595nm_ to monitor planktonic growth, and there was minimal effect by MMV688978 compared to the untreated control (Figure 4(b)). MMV675968, MMV688179, and MMV687747 did not reproduce the biofilm inhibition observed during the preliminary screening in the concentration-dependent assay.

**Figure 4.**
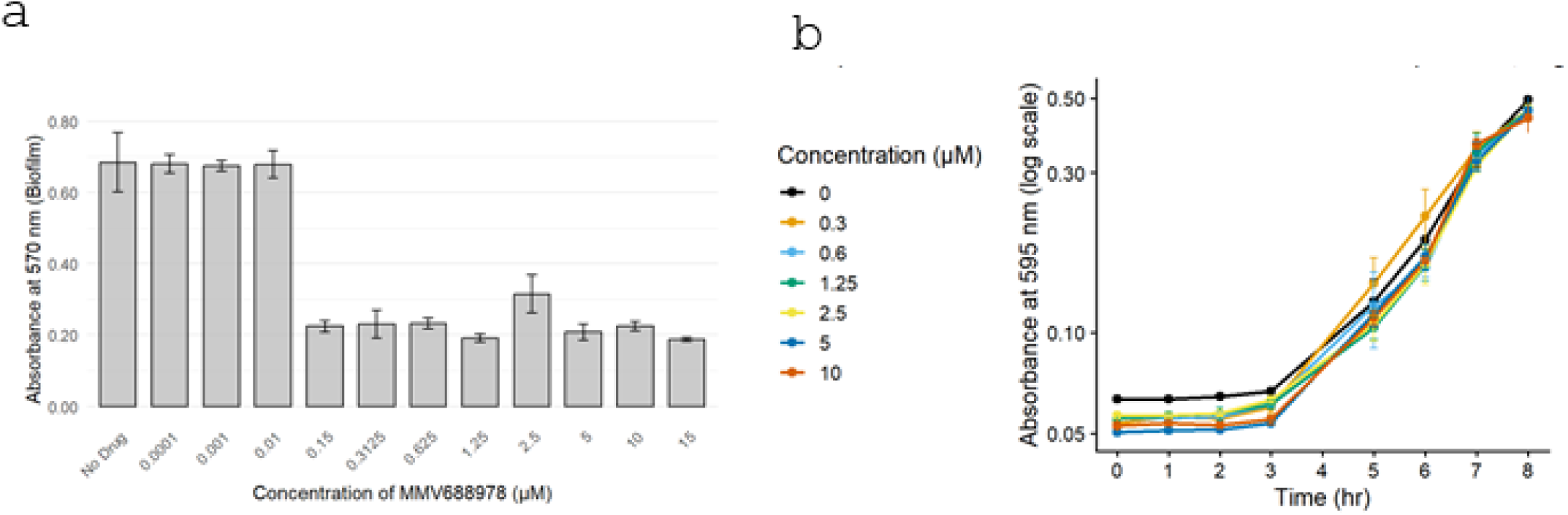
(a) MMV688978 inhibits *S*. Elisabethville LLD035X biofilm formation in a dose-dependent manner without affecting bacterial viability. (a) Quantification of biofilm formation at 24 h using crystal violet staining in the presence of increasing concentrations of MMV688978. Biofilm inhibition ranged from 49% to 72% across concentrations of 0.15–15 μM (P < 0.001) relative to solvent-treated controls; 0.0001–0.01 µM (P > 0.05). Data represent mean ± SE. Statistical significance was determined by one-way ANOVA followed by Dunnett’s multiple comparisons test versus solvent control (0 µM). (b) Growth of planktonic bacteria in the presence or absence of MMV688978 (5□µM) over 8□h. Absorbance at 595□nm was measured at the indicated time points (0, 1, 2, 3, 5, 6, 7, and 8 h). Data represent mean ± SD of three replicates. MMV688978 did not affect bacterial growth.

### MMV688978 demonstrates broad-spectrum anti-*Salmonella enterica* biofilm formation spectrum

To determine whether the biofilm-inhibiting activity of MMV688978 extends beyond the strain used for preliminary screening, the compound was tested against a panel of other biofilm-forming *Salmonella*, including typhoidal (n=6) and non-typhoidal (n=8) isolates. As shown in Figure 5a, biofilm formation by all tested Typhi strains was inhibited by 30% or more by MMV688978. A more variable picture was seen among the non-typhoidal *Salmonella* with inhibition above 30% seen for *Salmonella* I 21:b:-CHH008A (89.80%) and *S*. Riverside strains JKH041X and JKH42X (80.80% and 55.33%, respectively). Modest inhibition was seen for *S.* Give LLH074X (47.37%), *S.* Give LLH72X (3.00%) and *S.* Enteritidis LLH212X (27.40%) were not inhibited at levels equivalent to the hit criterion.

**Figure 5.**
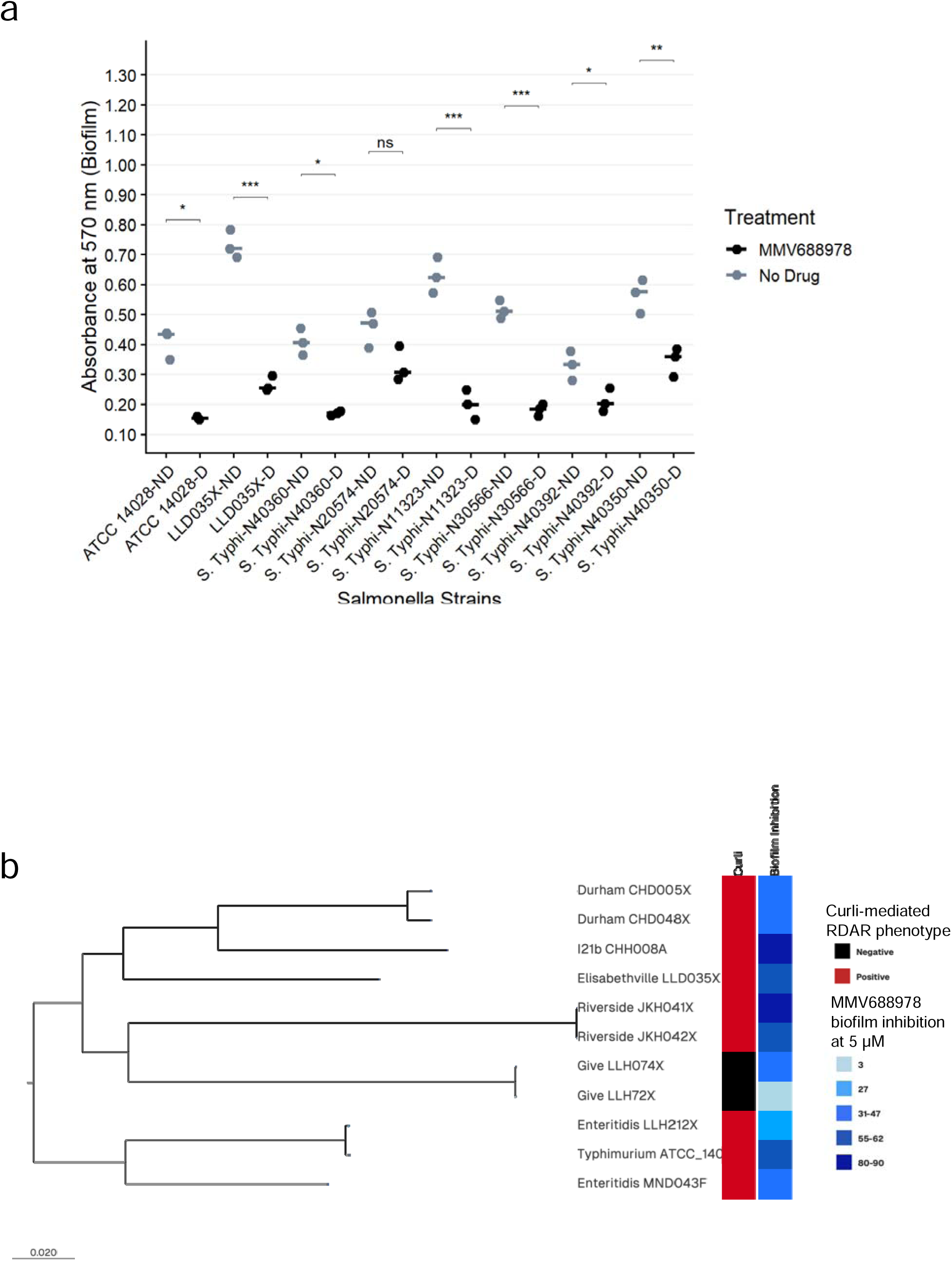
MMV688978 at 5 µM inhibit biofilm formation in both typhoidal *Salmonella* (NTS) and non-typhoidal isolates (a & b). (a) MMV688978 antibiofilm activity extends to *S*. Typhi from typhoid fever patients in Nigeria. Biofilm inhibition ranges from 66.93% for *S.* Typhi-N11323 to 28.68% for *S*. Typhi-N40350. Bars represent the median for replicates. (b) Phylogenetic distribution of auranofin-mediated biofilm inhibition illustrating the genetic relatedness of NTS isolates with phenotypic data for curli expression. MMV688978 (5 µM) inhibited biofilm formation of NTS expressing curli phenotype, inhibiting biofilm formation by 89.80% in CHH008A, 80.77% in JKH041X and 55.33% in JKH042X but inhibiting little above 40% in CHD005X, CHD048X, MND043F. Exception are *S.* Give LLH074X and *S.* Give LLH72X that does not express curli but whose biofilm formation was inhibited by 47.37% and 3.00%, respectively. Visualization was performed using Microreact to integrate phylogenetic clustering with antibiofilm phenotypes. MMV688978 demonstrated lineage-independent activity against curli-associated biofilm formation in NTS.

Beyond *Salmonella*, MMV688978 has been retrieved in previous antibiofilm screens against *Bacteroides fragilis* (Jang & Eom, 2020), *Staphylococcus aureus* and *Staphylococcus epidermidis* (Ferretti et al., 2025), *Aspergillus fumigatus* (Chen et al., 2023), EAEC 042 (Kwasi et al., 2022) suggesting that it may have an antibiofilm target that is not be *Salmonella*-specific. However, in this study, biofilm inhibition of *S*. Enteritidis LLH212X was less than 30%. It is unlikely (but not impossible) that a broad bacterial target is present in some *Salmonella enterica* serovars but not others, auranofin could target different factors in different species.

### MMV688978 interferes with motility but not curli

Surface proteins (e.g. outer membrane proteins) and appendages (fimbriae and flagella) are important for *Salmonella* biofilm formation. Below human body temperature, in environmental and reservoir niches, biofilm-forming *Salmonella* have been shown to produce a red, dry, and rough (RDAR) phenotype, which is detectable on Congo red agar and requires curli and cellulose production. All but one of the strains produced an RDAR phenotype *in vitro* (Figure 5b). Biofilm formation by the strain that did not exhibit this curli-dependent phenotype, *Salmonella* Give LLH074X was lower than median (55.33%) (Figure 5b) but still inhibited by MMV688978 (auranofin). Moreover, RDAR typically appears at stationary phase (Sjöbring et al., 1994; Ben Nasr et al., 1996; Römling et al., 1998), and our data show that, on *Salmonella*, auranofin inhibited biofilm formation at early but not late time points (Figure 6).

**Figure 6.**
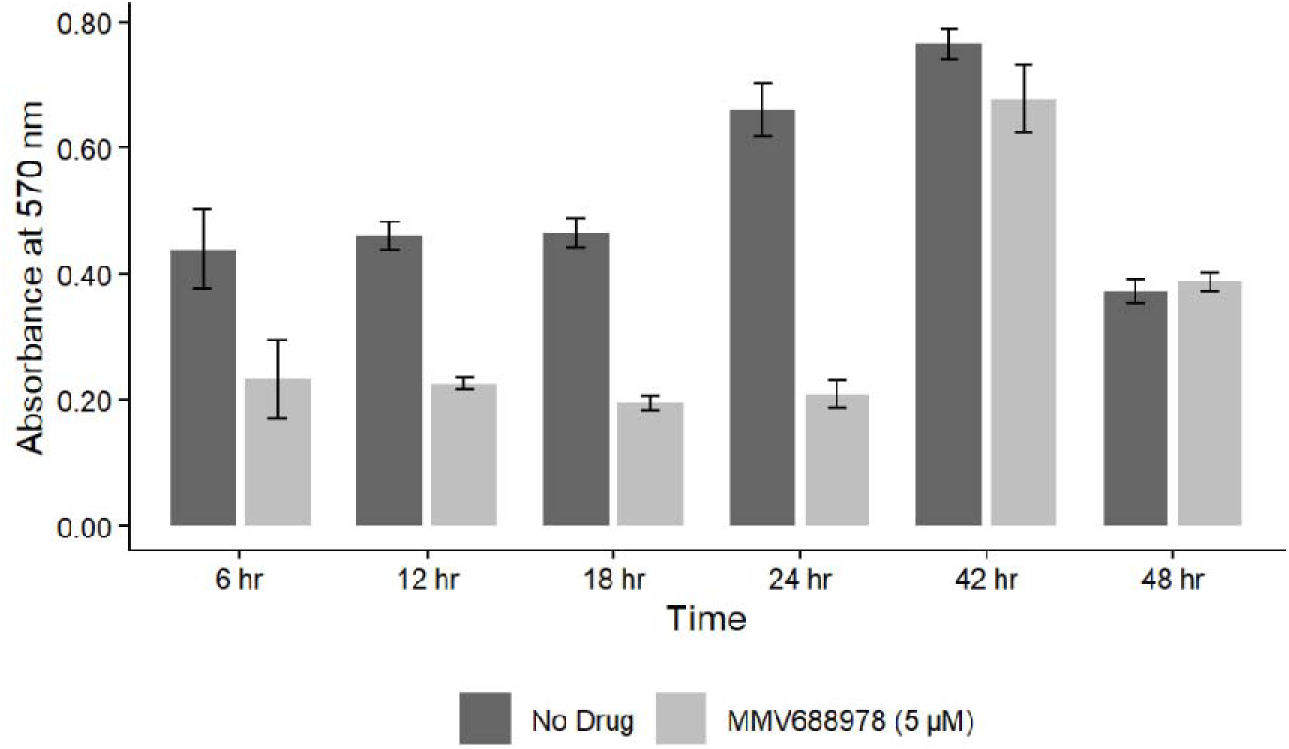
Biological target for MMV688978 may involve proteins responsible for early stage of *S*. Elisabethville LLD035X biofilm formation (a) MMV688978 demonstrated progressive antibiofilm activity 47.08% at 6 h to above 60% at 24 h (p < 0.001), after which activity declined. This suggest that the likely target for MMV688978 is involved in the initial phase of biofilm formation such as attachment and microcolony formation. Error bars represent the range of replicates.

Early-stage biofilm formation in *Salmonella* requires motility as well as adhesins (Berne et al., 2015; Crawford et al., 2010; Toguchi et al., 2000), and their regulators. We therefore evaluated the effect of MMV688978 effect on motility in sloppy agar. Auranofin inhibited visible motility of *S*. Elisabethville LLD035X at concentrations of 5 µM and 10 µM, but not at lower concentrations. As all *Salmonella* strains used in this study were motile we employed a non-motile Enterobacterales strain, *K. pneumoniae* ATCC 700603 to test the hypothesis that auranofin might be interfering with motility, As illustrated in Figure 7, MMV688978 exhibited only weak antibiofilm activity against this bacterium, with an inhibition of only 6.53% at 5 µM, compared with *S.* Elisabethville LLD035X, *S*. Typhimurium ATCC 14028 at the same concentration for motile *Salmonella* strains.

**Figure 7.**
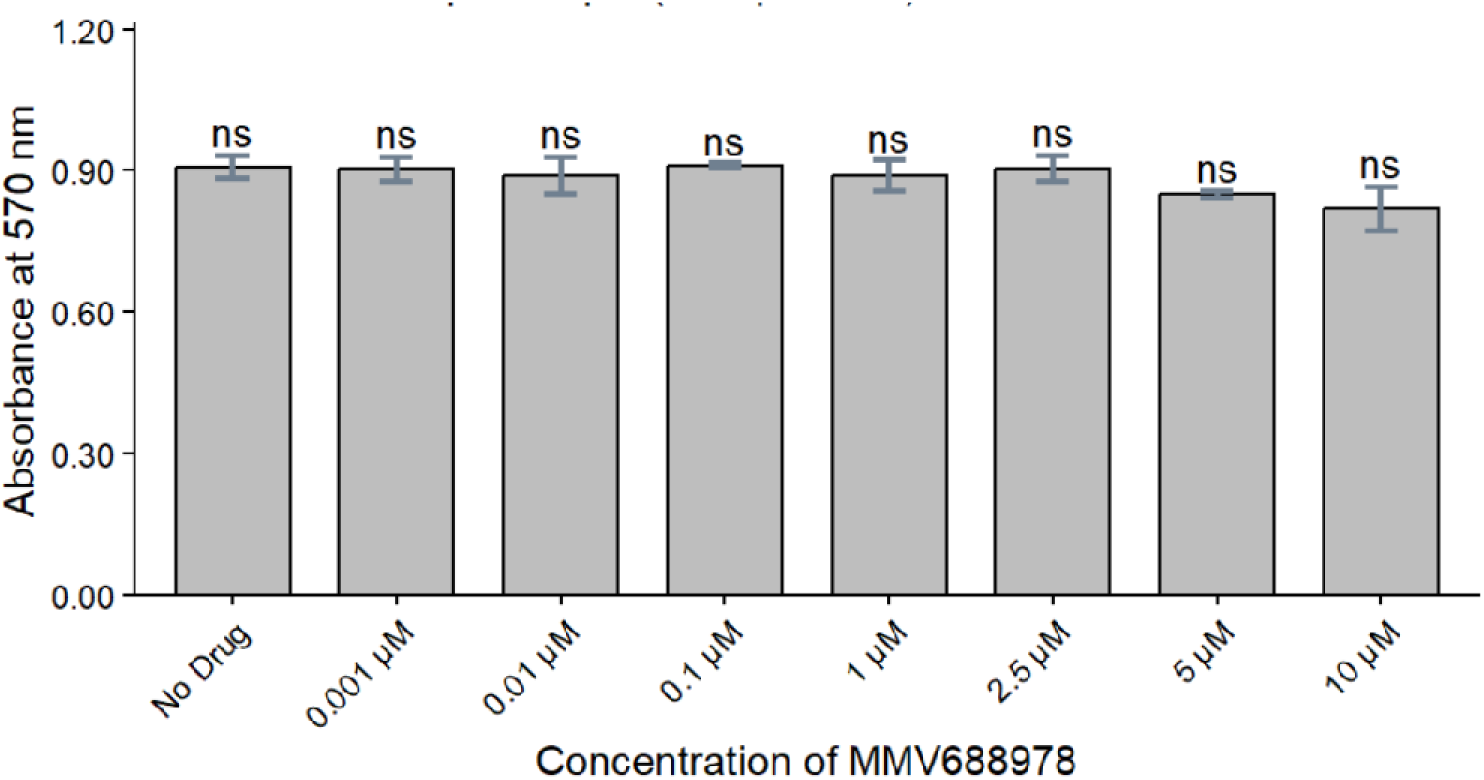
MMV688978 inhibits biofilm formation of a non-motile bacterium, “*K. pneumoniae* ATCC 700603” by 6.53% and 8.70% at 5 μM and 10 μM, respectively, an effect lower than what is seen in the inhibition of *S*. Elisabethville LLD035X biofilm formation. The low activity of MMV688978 as an antibiofilm agent against a non-motile bacteria suggest the activity of the compound may involve interference with the motility pathway.

### Auranofin’s gold (I) ion is essential for its *Salmonella* antibiofilm activity

**The** Auranofin molecule consists of a gold (I) ion linked to triethylphosphine and attached to a tetraacetylated thiosugar (a). An analogue of MMV688978, aurothioglucose (b), which retains the gold ion but lacks many of the auraofin side chains, and a non-gold analogue of MMV688978, 1-thio-beta-D-glucose tetraacetate, which retains the side-chain (c), were tested for their antibiofilm activity against *S*. Elisabethville LLD035X. Quantification of eluted crystal violet from 24-hour biofilms revealed that at 5 μM, aurothioglucose inhibited *S*. Elisabethville LLD035X biofilm formation by 61.30%, comparable to auranofin and significantly higher(P < 0.001) than non-gold compound, 1-Thio-beta-D-glucose tetraacetate (Figure 8 a-e). This suggests that gold (I) is vital for the antibiofilm activity of MMV688978. We then determined whether auranofin and its derivatives had any effect on *Salmonella* motility. At 5 µM, auranofin and aurothioglucose inhibited the motility of *S.* Elisabethville LLD035X.

**Figure 8.**
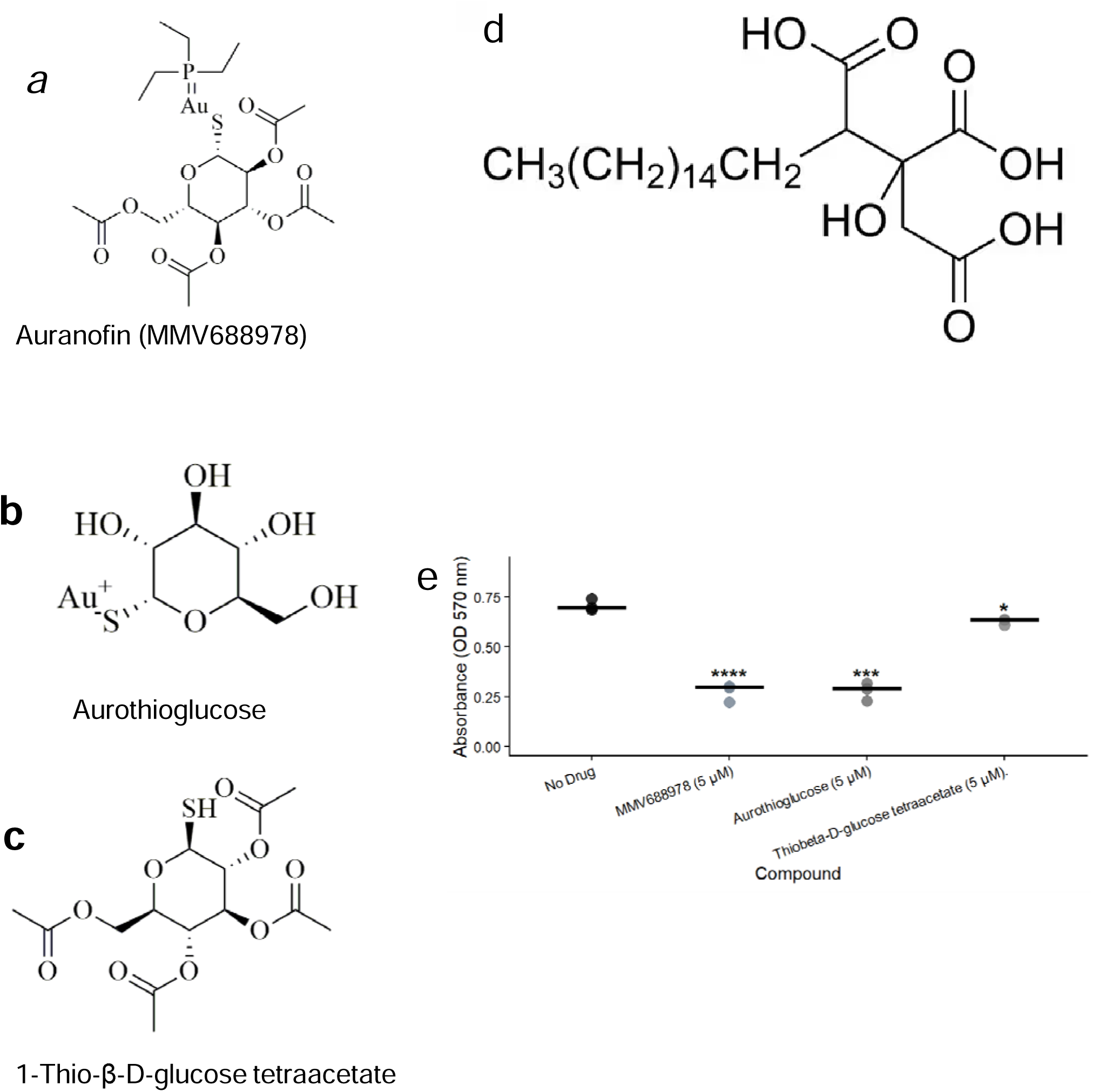
Aurothioglucose, a gold analogue of auranofin, inhibits *S*. Elisabethville LLD035X biofilm formation (a-d). (a) Chemical structure of MMV688978 (auranofin). (b) Chemical structure of aurothioglucose (c) Chemical structure of 1-thio-β-D-glucose tetraacetate (d) Chemical structure of agaric acid and (e) Gold analogues such as auranofin and aurothioglucose inhibit biofilm formation of *S*. Elisabethville LLD035X above 60% in contrast to a non-gold component of auranofin, 1-thio-β-D-glucose which demonstrated 11.39%. Bars represent the median of three replicates.

## Discussion

Formation of biofilm by *Salmonella* strains is known to be central to their pathogenesis and transmission (Pradhan et al., 2022). *S. enterica* biofilm formation is regulated by a tightly controlled mechanistic cascade that involves adherence by fimbrial and surface factors, flagellar motility, production of cellulose and other matrix components, which are regulated by specific transcriptional regulators (Gerstel & Römling, 2003; Steenackers et al., 2012). Targeting biofilms is widely reported to impose less evolutionary pressure than known bacteriostatic/bactericidal agents, which select resistant mutants (Wang et al., 2021; Roy et al., 2018).

To date, antibiofilm agents with activity against *Salmonella enterica* have not been reported for clinical use or testing, although primary investigations have been conducted, particularly in the last decade. Studies that have investigated antibiofilm activity of small molecules from both of natural and synthetic sources against *Salmonella* strains, typically evaluated a single molecule, structurally related molecules or molecules with a similar mode or mechanism of action (Lories et al., 2020; Moshiri et al., 2018; Huggins et al., 2018; Dias de Emery et al., 2023; Vidaković Knežević et al., 2025; Kim et al., 2022; Sandala et al., 2020; Merino et al., 2025). In contrast, this study deployed a curated library of 400, mostly unrelated drug-like compounds implicated for different indications (MMV, 2019). We hypothesized that an unbiased chemical screen would allow us to identify hits with vastly different mechanisms of action and the potential to repurpose existing compounds if highly active antibiofim agents were identified. Moreover, performing the screen with a small library, the MMV Pathogen Box, which gave a hit rate of 0.75% provides proof of concept, justifying future screens.

We applied hit-selection criteria of ≥30% biofilm inhibition and ≤10% growth inhibition, which have been used in other studies (Kwasi et al., 2022; Koopman et al., 2015). The low hit rate is comparable to those from other studies involving biofilm-forming pathogens such as *Candida albicans* (Vila and Lopez-Ribot, 2016), and enteroaggregative *Escherichia coli* (Kwasi et al., 2022), as well as *S. enterica* (Koopman et al., 2015), all of which yielded hit rates of <2%. Thus, antibiofilm activity is not a nonspecific, trivial, or incidental phenotype and likely but requires definite perturbation of specific targets and/or regulatory networks.

Biofilm-forming mechanisms differ among *Salmonella* serovars and even between different strains from the same lineage. It was therefore important for us to screen more than one strain and the lack of overlap in hits, as well as the differences in activity of hits among different strains, suggests that broader strain panels should be used in future screens.

The screen yielded MMV688371 as a hit against *S*. Typhimurium ATCC 14028, as well as MMV688978 (auranofin), and MMV687273 (ethambutol analogue) as hits against *S*. Elisabethville LLD035X. These compounds inhibited biofilm formation by > 30% while maintaining growth inhibition below 10%. MMV687273 (ethambutol analogue) and MMV688371 (a benzamide) have previously demonstrated antimalarial activity; with MMV688371 demonstrated activity against *Plasmodium falciparum* with an IC_50_ of 5.85 µM (Patra et al., 2019). MMV687273 displayed activity against *P. falciparum*, exhibiting an IC_50_ of 0.105 µM for stage IV/V gametocytes (Reader et al., 2021). Both hits were also reported to be effective against *Trypanosoma species*. MMV688371 demonstrated an IC□□ of 1.60□µM against *Trypanosoma brucei brucei* (Dize et al., 2022) and, the activity of MMV687273 resulted in a mortality rate of 75.8% of *Trypanosoma evansi* (Canever & Miletti, 2020). In addition, MMV687273 inhibits *Fonsecaea pedrosoi* growth by 37% (Coelho et al., 2020) and was found to have synergistic effect when combined with itraconazole against this pathogen (Coelho et al., 2022). MMV687273 has previously shown to demonstrate antibiofilm activity against *C. albicans* (Vila and Lopez-Ribot 2017) and could potentially have a broad spectrum antibiofilm target. After the preliminary screening, we could not proceed with the validation of MMV688371 and MMV687273 because they were not available but there are plans to obtain them for further experiments.

MMV688978 (auranofin) is an antirheumatic drug but of recent has been the subject of anticancer investigations (Abdalbari and Telleria, 2021). It has also shown to have activity against *Mycobacterium tuberculosis* (Harbut et al., 2015) and *Helicobacter pylori* (Epstein et al., 2019). However, it does not have known anti-Gram negative activity, beyond *H. pylori,* due to its inability to permeate the outer membrane (Cassetta et al., 2014; Harbut et al., 2015; Hokai et al., 2014; Thangamani et al., 2016). In this study, we recorded growth inhibition of *Salmonella* screened as −14.45% for *S.* Elisabethville LLD035X, - 12.93% for *S.* Riverside JKH041X and −14.50% for *S.* Riverside JKH042X, showing that the organisms were able to grow slightly better in the presence of the drug. In contrast we found that it inhibited the growth of *S.* Typhimurium ATCC 14028 by 17.80%. Thus, the compound does inhibit growth of some strains but shows robust biofilm inhibition in resistant strains, pointing to biofilm-specific target(s). MMV688978 demonstrated activity against typhoidal and non-typhoidal *Salmonella* serovars. This implies that its targets may be conserved across the subspecies, although expression or accessibility of the target may vary among *Salmonella enterica* strains. *Salmonella* serovars are known for their heterogeneity and genetically diverse in biofilm regulation and formation (Beshiru et al., 2018, Zhang et al., 2025). Cross-serovar activity is important due to the wide variety of lineages that cause human disease and their distribution in a range of niches, with different transmission pathways. The spectrum of the drug is broader than Salmonella: previously, MMV688978 was found to show antibiofilm activity against EAEC (Kwasi et al., 2022), *Staphylococcus aureus* 700699 (Bhandari et al., 2018) and C*andida albicans* (Siles et al., 2013). In addition to pointing to an unusually broad spectrum for an antibiofilm agent, these findings imply that MMV688978 could interfere with fundamental processes involved in biofilm formation.

*Salmonella* and other Enterobacterales form biofilms by initial attachment and motility, autoaggregation to form clusters and building a macromolecular matrix (Koopman et al., 2015). The specific factors that mediate these functions and the regulatory systems that control them differ among species, subspecies and even strains. However, factors such as chaperone-usher family pili, flagella and curli are common to many different species, and are used under different conditions. A number of different biofilm-forming conditions for *Salmonella* are documented in the literature. We found that most strains in our collection formed good biofilms in 0.5X TSB at 28°C, conditions that mimic environmental niches that *Salmonella* may encounter in livestock and the foodchain as well as in wastewater, therefore determining transmission success (Schonewille et al., 2012; Marin et al., 2009). MMV688978 demonstrated a progressive biofilm inhibition up to 24 h, afterward its activity declined. This implies that its activity affects early stages of biofilm formation, such as attachment to surfaces. Preventing the attachment phase hinders the establishment of community-embedded biofilm architecture (O’Toole et al., 2000) and such compounds with early activity could serve as adjunctive therapy during infection (Zhou et al., 2025), as well as colonization of environmental niches. The diminished activity of MMV688978 after 24 h suggests that it may lack accessibility to the internal environment of biofilm, once the exopolymer matrix has matured or less relevance of its target in late stages (Venkatesan et al., 2015; Hall and Mah, 2017; Fu et al., 2021). This observation agrees with the finding that MMV688978 did not impact the curli- and cellulose-mediated RDAR late-stage phenotype but did impact motility.

At concentrations of 5.0 µM and 10.0 µM MMV688978 (auranofin) inhibited swimming motility of *S*. Elisabethville LLD035X. In motile Enterobacterales, such as *Salmonella*, flagellar motility is a prerequisite for surface attachment (Crawford et al., 2010; Haiko & Westerlund-Wikström, 2013). In the absence of flagella, planktonic cells may not be able to reach suitable surfaces and initiate attachment due to the repulsive physicochemical forces, thus hindering biofilm development. Chemical agents such as agaric acid (Lories et al., 2020) and brominated furanones (Janssens et al., 2008) have been reported to inhibit *Salmonella* biofilm formation by interfering with motility. Agaric acid shares some structural similarity with auranofin and inhibits early-stage biofilm formation, without affecting growth, by reducing expression of class II and III flagella genes. Agaric acid is also active against *E. coli* biofilms but does not inhibit the formation of biofilms by non-motile *Staphylococcus aureus* (Lories et al., 2020). Like agaric acid, and brominated furanones, auranofin demonstrated a broad spectrum and motility-inhibiting activity. Auranofin inhibits biofilm formation of both enteroaggregative *E. coli* (Kwasi et al 2022) and a range of *Salmonella* isolates in this study but did not inhibit biofilm formation by non-motile *Klebsiella* strain ATCC 700603. Auranofin (Molecular weight = 678.48 g/mol) is larger than brominated furanones (molecular weight range = 140.18 to 422.22 g/mol) and agaric acid (molecular weight = 416.55 g/mol), does not penetrate the cell to exhibit antibacterial activity, and so, despite minor structural similarities to agaric acid, unlikely to have an intracellular motility target. While we did not have the mutants and constructs required to identify its precise target, by comparing the activity of the auranofin molecule to its analogues aurothioglucose and 1-thiobeta-D-glucose tetraacetatex, we determined that the gold (1) moiety is required for activity and the auranofin side chains that resemble agaric acid are not. The apparent requirement for gold is a negative, since most of the adverse effects of auranofin and aurothioglucose are attributable to their gold content. However, although lead progression of auranofin for *Salmonella* can be little justified at this point, understanding auranofin’s target may open the possibility of identifying other inhibitors.

Some of the limitations of the study are that we did not screen for compounds that can inhibit *Salmonella* biofilm formation under conditions analogous to those *in vivo*, nor did we screen a broad range of strains initially. However, the discovery of MMV688978 as an antibiofilm agent with broad activity spectra against *Salmonella enterica* underscores the possibility of targeting virulence and highlights the usefulness of drug repurposing in drug discovery project. Further work on MMV688978 will identify the specific molecular target that accounts for its antibiofilm and antimotility activity, optimise it and evaluate its utility for prevention of Salmonella transmission or infection alone or in combination with antibacterials. Future studies will also need to evaluate and characterize the hits MMV687273 and MMV688371 as well as identify the specific target of auranofin (MMV888978).

In conclusion, this study identified drug-like, antibiofilm agents from MMV Pathogen Box. Auranofin, in particular, is a potent, concentration-dependent inhibitor of S. Elisabethville LLD035X biofilm formation with activity extending to other serovars of *Salmonella*. Its inhibitory activity is dependent on the gold (I) ion of the molecule, limiting potential lead for therapeutic development. The study, like that of Lories et al (2020) and Janssens et al (2008) has also uncovered motility as an important antibiofilm target, for which specific screens could be developed to identify other antibiofilm scaffolds.

## Acknowledgements

The following people are appreciated for the technical assistance, logistical support and useful comments offered during the course of this study: Stella E. Ekpo, Oyeniyi Stephen Bejide, Rotimi Dada, Mariam A Odebode, Jola-Ade J. Ajiboye and other staff and students of the Molecular Microbiology Research Laboratory, Faculty of Pharmacy, University of Ibadan, Nigeria.

